# Change-resistance distinguishes the representational geometries of human spatial memory and mouse CA1 in deformed environments

**DOI:** 10.1101/2025.07.09.663962

**Authors:** Diana M. Green, David G. Behery, Rachel E. Barrett, Viktor H. Klimczyk, Michael Pigin, Sergio A. Pecirno, Alexandra T. Keinath

## Abstract

Prior work has highlighted qualitative similarities between the neural instantiations of cognitive maps in rodents and memory-guided navigation in humans, suggesting a conservation of representational structure across species. Yet evidence of cross-species differences in neural coding continues to mount. Our ability to reconcile these similarities and differences has been inherently limited by the qualitative nature of our cross-species comparisons. To overcome this limitation, here we combine recent technical and theoretical advances to characterize the representational geometry of human spatial memory during a diverse set of environmental deformations and compare this geometry to that of mouse CA1. Across three untethered immersive virtual reality experiments (n > 100 participants per experiment), we find that deformations induce compounding local distortions in human spatial memory. These distortions yield a representational geometry which closely resembles a change-resistant version of that of mouse hippocampal CA1 during analogous deformations. The geometries of mouse CA1 subpopulations with higher firing rates, spatial tuning stability, and spatial tuning specificity all better resembled that of human spatial memory. The precision, but not accuracy, of human spatial memory also modulated cross-species resemblance. The local impact of deformations scaled up when humans navigated a larger environment, preserving representational geometry and cross-species resemblance. Neither geometry nor cross-species resemblance depended on the human visual advantage during retrieval. Together, these results establish a common cross-species resemblance in the representational geometry of mouse CA1 and human spatial memory during environmental deformations, with a notable difference in the resistance to change between these assays.

---

Organisms from mice to humans rely on mnemonic representations of the world and their relationships to it to flexibly and efficiently navigate our dynamic world^1^. Referred to as *cognitive maps*^2^, these representations are instantiated by the coordinated activity of neural populations throughout the brain tuned to navigationally relevant content. These include *place cells* representing contextualized location in CA1 and CA3 of the hippocampus^3,4^, as well as extra-hippocampal populations representing locations^5–7^, distances^6,8^, directions^9–12^, goals^13–15^, objects^16–19^, and boundaries^20–23^.

Extensive work has investigated the determinants of cognitive mapping in both rodents and humans by assaying its neural and behavioral correlates while manipulating features of the world^7,24–31^. In many cases, this work has highlighted qualitative similarities between the impacts of environmental manipulations on the neural representations of rodents and the spatial memory of humans. For example, when a familiar environment is reshaped by stretching or compressing one or both of its dimensions, navigationally relevant neural codes in rodents – including the hippocampal spatial code – rescale to roughly match the environmental deformation^7,27,28^. Similarly, when human participants are asked to replace objects at remembered locations in a rescaled environment, their judgements will roughly scale in concert with the environment^25,31,32^. These and other similarities^26,29,33^ provide provocative evidence that cognitive maps are evolutionarily conserved representations which might be instantiated by common cross-species mechanisms. On the other hand, mounting work has led to a growing appreciation for the differences between the neural instantiations of cognitive maps across species, especially differences in temporal structure^34–37^ and the primacy of visual correlates in highly visual animals such as primates and certain birds^38–45^.

While these comparisons have been foundational to building a cross-species understanding of cognitive maps, reconciling similarities and nuanced differences will require us moving beyond qualitative similarity toward a more precise, quantitative comparison of representational structure across species and assays. Though a challenge for traditional experimental approaches^46^, recent complementary technical and theoretical advances have laid the groundwork to make such comparisons possible^47^. This theoretical work has emphasized how the geometry of a representation is determined by its neural instantiations and informs its contributions to cognition^48,49^. Leveraging modern behavioral and neural recording techniques, this *representational geometry* is something that can be characterized with sufficiently high precision to be quantitatively compared across assays, even if those assays are derived from different recording techniques or species. Doing so will help us move beyond qualitative comparisons and come to more confident conclusions about which neurocognitive mechanisms are conserved across species and which are not.

Motivated by this approach, here we characterize the representational geometry of human spatial memory during a diverse set of environmental deformations and quantitatively compare this geometry to that of mouse hippocampal CA1. To assay human spatial memory, we leveraged untethered immersive virtual reality (VR) in large cohorts of participants (n > 100 per experiment) tasked with remembering and replacing objects in parametrically deformed versions of a familiar environment. Across three experiments, we find that deformations induce compounding, local distortions in human spatial memory. These distortions yield a representational geometry which closely resembles a change-resistant version of that of mouse hippocampal CA1 during analogous deformations. The geometry of mouse CA1 subpopulations with higher firing rates, spatial tuning specificity, and spatial tuning stability all better resembled that of human spatial memory. The precision, but not accuracy, of human spatial memory also modulated cross-species resemblance. Representational geometry and cross-species similarity did not depend on the availability of distal visual cues during retrieval or the scale of the environment that human participants navigated. Together, these results establish a common cross-species resemblance in the representational geometry of mouse CA1 and human spatial memory during environmental deformations, with a notable difference in the resistance to change between these assays.

## Results

### Assaying human spatial memory in deformed environments using immersive VR

We leveraged untethered immersive VR to characterize the impact of a diverse set of environmental deformations on human spatial memory. To this end, we taught participants (n = 104) the locations of 5 objects pseudo-randomly distributed in a 5 x 5 m open square space and later tested their memory for these objects in the square space as well as nine deformed configurations (Fig. 1a,b). Each deformed configuration was created by imagining the square environment as a 3 x 3 grid of partitions and closing combinations of these partitions. The configurations were chosen to match those previously used to characterize mouse hippocampal CA1^50^ to enable quantitative comparison of the impact of deformations on representational geometry across species, as described below.

**Figure 1.**
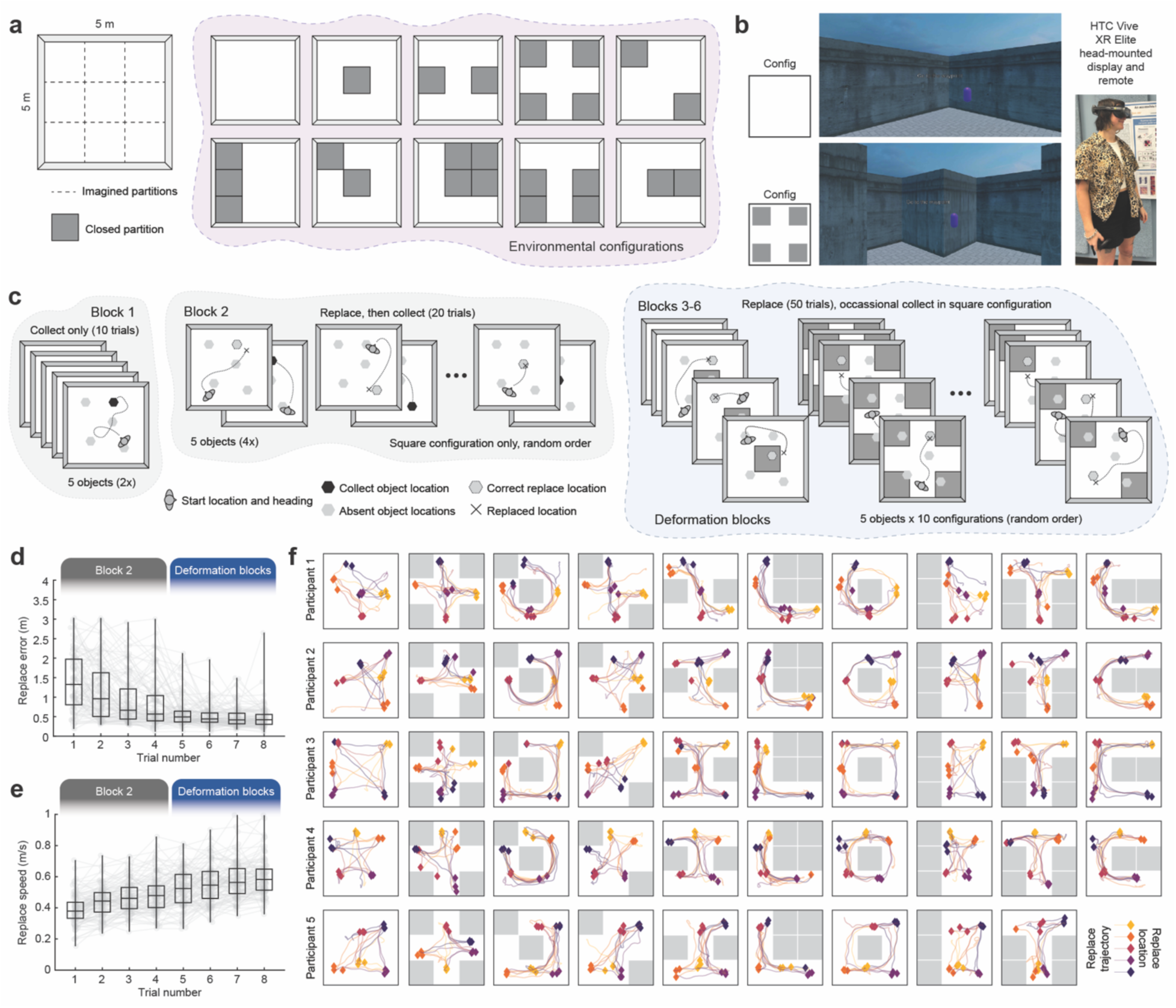
(**a**) Schematic of all ten environmental configurations in which human spatial memory was tested. (**b**) Example views of the environment in configurations with and without closed partitions as well as a picture of a participant completing the experiment. (**c**) Schematic of experimental design. (**d**) Error in replace locations in the square configuration across blocks. (**e**) Replacement speed in the square configuration across blocks. (**f**) Example replacement trajectories and locations for all objects during all deformation block trials for five example participants.

The experiment followed a blocked design motivated by prior work^31,51^, with participants repeatedly collecting and replacing each object one by one to learn and assess their memory (Fig. 1c). In the first block, the participant collected each object twice in a random order. In the second block, the participant tested their memory by replacing each object four times in a random order in the square configuration. After each replacement, the participant was given an opportunity to collect the object from a new starting point to improve their memory. Finally, in blocks three through six the participant replaced each object in each configuration (including the square configuration) once per block in a random order. We refer to these blocks as ‘deformation blocks’. Because the objects were only ever collected in the square configuration, there was no ‘correct location’ to replace the object in the deformed configurations. Instead, the participants were instructed to respect the virtual boundaries and replace each object at the location they think best. During each deformation block the participant was also given three more opportunities to collect each object in the square configuration to continue improving their memory. Before collect trials and between configuration changes, the participant was guided to a new location by a waypoint whose location they did not need to remember.

### Environmental deformations locally deform human spatial memory

Over the course of the experiment, participants moved more quickly and became more accurate in their replace locations in the open square configuration (Fig. 1d,e). In deformed configurations, participants made qualitatively sensible decisions about where to replace objects, despite the lack of a ‘correct’ location (Fig. 1f). Qualitatively, these decisions sometimes included varying the replacement locations for the same object, replacing the same object on different sides of closed partitions, and changing the replace locations of some objects but not others.

To quantify these observations, for each object and configuration we computed the mean distance of all four replacements to the median replace location in the square configuration, a measure we refer to as the *replacement spread* (Fig. 2a). If deformations have an impact on human spatial memory, then we would expect the replacement spread of objects in deformed configurations to be greater than the replacement spread in the square configuration. Importantly, in some cases an object’s median replace location in the square configuration was no longer accessible during a given deformation. In those cases, an increase in replacement spread is guaranteed. Therefore, we focused our analysis only on the replacement spread for objects which remained accessible during deformations (Fig. 2a).

**Fig. 2.**
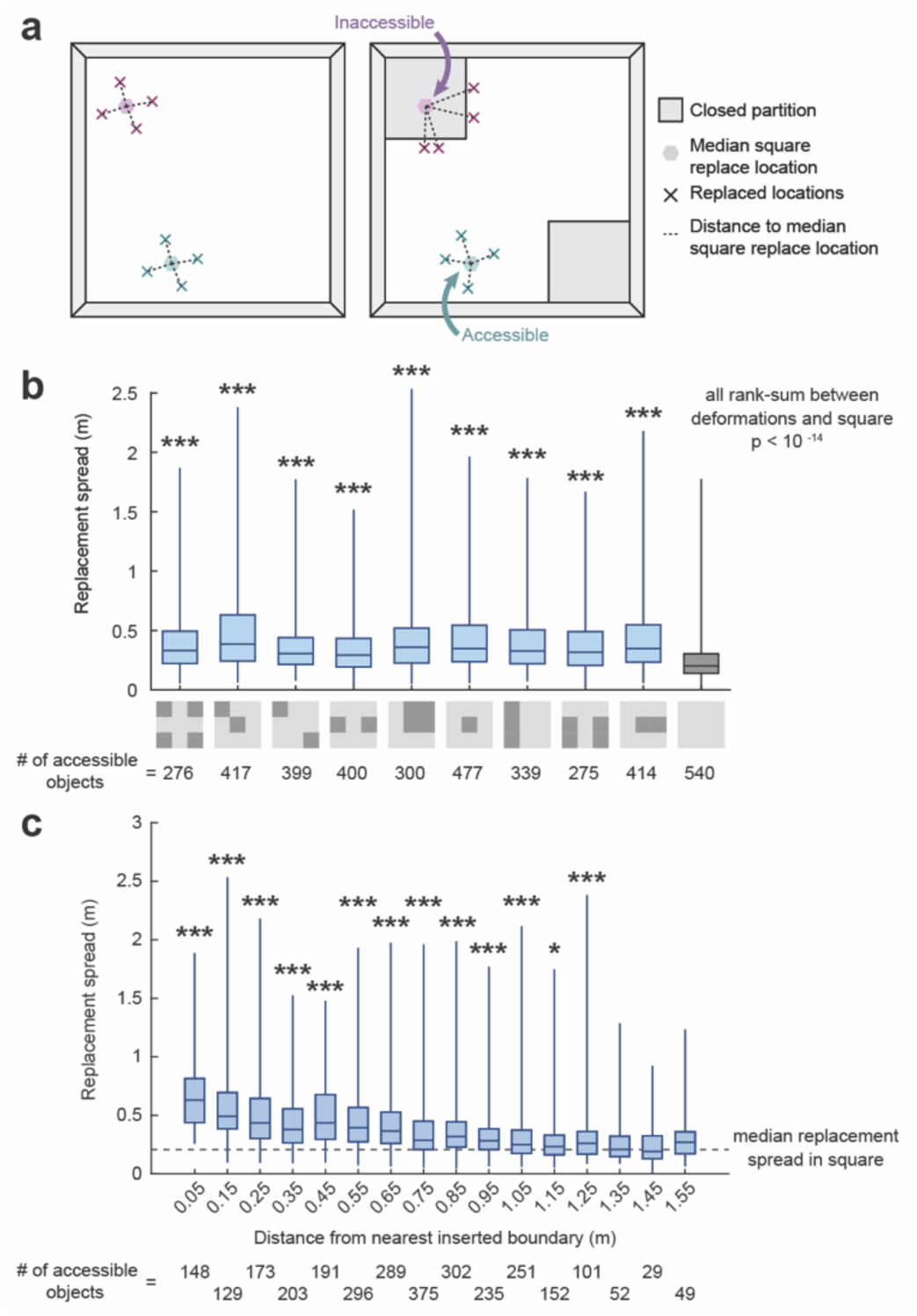
(**a**) Schematic of replacement spread analysis. Replacement spread was computed as the mean distance to the median square replace location within each configuration. Only objects with accessible median square replace locations were included in further analysis. (b) Replacement spread for each configuration. Significance markers indicate the outcome of a rank-sum test between the replacement spread of the denoted configuration and the square configuration. (**c**) Replacement spread as a function of distance to the nearest inserted boundary, aggregated across configurations. Significance markers indicate the outcome of a rank-sum test between the replacement spread of the denoted distance group and the square configuration. See Supplementary Figure 1 for unbinned data. *p<0.05, ***p<0.001, uncorrected

In all deformed configurations, we observed replacement spreads that were significantly larger than those measured in the square configuration (Fig. 2b). Note that this did not have to be the case, as only objects with accessible median square replace locations were included in this analysis. Next, we asked whether the increase in replacement spread depended on the distance from inserted boundaries. In rodents, environmental deformations induce local distortions in their spatial neural codes whose impact decreases as a function of distance from altered boundaries^30^. Aggregating accessible objects across deformed configurations, we found that replacement spread was greatest for objects with median square replacement locations near inserted boundaries (Fig. 2c). Replacement spread decreased as a function of distance to the nearest inserted boundaries but remained significantly elevated relative to that of the square configuration for objects replaced within 1.25 m of the nearest inserted boundary. These results demonstrate that environmental deformations have a significant local impact on human spatial memory, parallelling the impact of environmental deformations on rodent neural codes.

### Characterizing representational geometry in deformed environments at different resolutions

To better understand the impact of environmental deformations on human spatial memory, we next characterized the similarities and dissimilarities among patterns of replacements across configurations. To this end, we first transformed the replace locations of each object into a replacement map by binning and smoothing all four replacements in each configuration (Fig. 3a). Next, for each object we computed replacement map correlations between all pairwise comparisons of configurations and took the mean across objects (Fig. 3b). This yields a representational similarity matrix (RSM) which summarizes the overall similarity of replacement patterns between configurations (RSM; Fig. 3b). We refer to the similarity structure captured by the RSM as the *representational geometry* of that measure.

**Figure 3.**
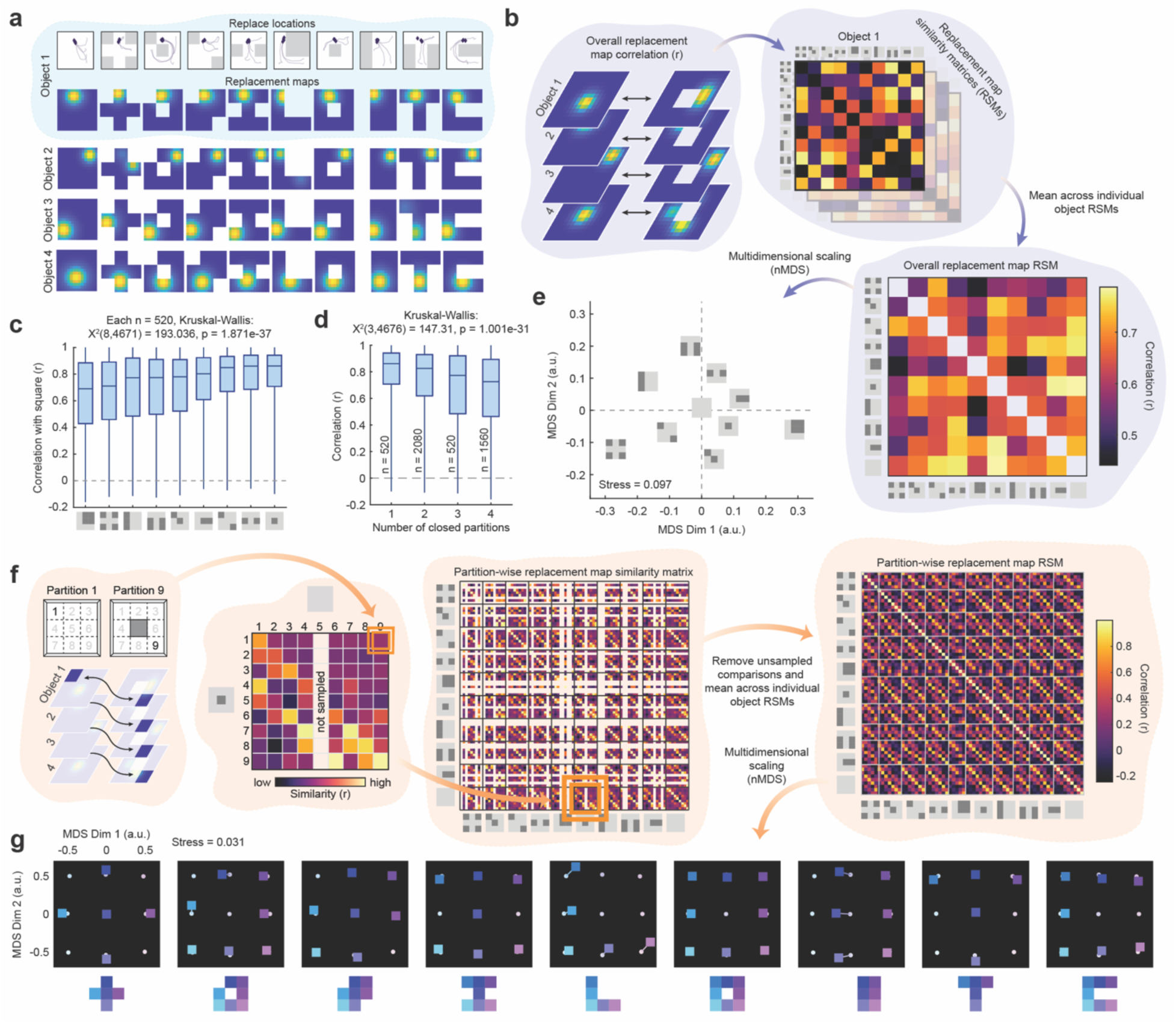
(**a**) Example replacement trajectories and replacement maps for four objects in all ten configurations. (**b**) Schematic illustrating the construction of overall replacement map representational similarity matrices (oRSMs). (**c**) Replacement map correlations between each deformed configuration and the square configuration. (**d**) Replacement map correlations between the deformed configurations and the square configuration as a function of the number of inaccessible partitions. (**e**) Embedding of configuration similarity derived from oRSM via nonmetric multidimensional scaling. Note that configurations which share closed partitions tend to be more similar, i.e. closer to one another in this embedding. (**f**) Schematic illustrating the construction of partition-wise replacement map representational similarity matrices (pRSM). (**g**) Embedding of partition similarity derived from pRSM via nonmetric multidimensional scaling. White dots indicate the embeddings of each partition in the square, while the colored boxes represent the embedding of the same partition in the deformed configuration. Note that closing partitions results in nearby partitions becoming more similar to the closed partition’s representation, reflect in the movement of nearby partitions toward closed partitions during the deformations.

To make sense of this representational geometry, we began by focusing on comparisons between the deformed configurations and the square configuration (Fig. 3c). We first noted that different deformed configurations varied in their similarity to the square (Fig. 3c), with similarity to the square generally decreasing as a function of the number of closed partitions (Fig. 3d). To visualize the full geometry, we performed nonclassical multidimensional scaling. Multidimensional scaling is a class of techniques for transforming distance matrices (i.e. 1 - RSM) into a low-dimensional embedding where the relative locations of points reflect the similarity of those points to one another as best as possible. In other words, multidimensional scaling recovers an approximation of the representational geometry which gave rise to the corresponding distance matrix. Nonclassical multidimensional scaling focuses on recapitulating the rank order distances rather than the raw distance values themselves and is particularly apt when working with correlation distances^52^. This visualization revealed that configurations with more closed partitions tended to be most dissimilar from other configurations, including one another (Fig. 3e). Deformed configurations which shared closed partitions also tended to be more similar to one another than configurations which did not share closed partitions. Altogether, these results demonstrate that deformations induce distinct and compounding distortions to human spatial memory.

In the previous analysis, we characterized the representational geometry of human spatial memory at the resolution of the entire environment. However, our parametric partition-based design also allows us to characterize representational geometry at a finer resolution. To do so, we computed the replacement map correlations between pairwise comparisons of partitions within and across all configurations (Fig. 3f). Because some partitions were inaccessible in some configurations, comparisons with those partitions were excluded. This RSM characterizes the impact of deformations on the representational geometry of human spatial memory at a finer resolution than that of the overall replacement map RSM, complementing our previous analysis. To distinguish the two, we refer to the overall replacement map RSM as the oRSM and the partition-wise RSM as the pRSM.

To make sense of the pRSM, we again used nonmetric multidimensional scaling. Comparing embeddings of the same partitions between the deformed configurations and the square configuration, we found that deformations induced modest distortions with interpretable characteristics (Fig. 3g). For example, closing a partiton tended to make the representations of nearby partitions more similar to the closed partition, represented by movement of the nearby partitions toward the closed partition in the embedding (Fig. 3g; note configurations L, T, C, and rectangle). This pattern was particularly striking in the rectangular configuration, where only the partitions neighboring the closed partitions were impacted. These results are consistent with a local impact of deformations and contrasts with uniform rescaling predicted by some theories^7,24,27^. Altogether, these findings demonstrate that deformations induce modest but interpretable distortions in the fine-resolution geometry of human spatial memory.

### Comparing representational geometry in deformed configurations across species

In the prior section, we characterized the representational geometry of human spatial memory in deformed environments at two resolutions. While this characterization is informative on its own, an additional advantage of this approach is that it enables quantitative comparison with the representational geometry of other assays, even if they are recorded using other techniques or are derived from other species^47,48^. In prior work from the Brandon lab^50^, we characterized the representational geometry of hippocampal CA1 in mice exploring analogously deformed environments. Leveraging miniscope imaging^53^, we simultaneously recorded large populations of neuron from hippocampal CA1 as mice explored each of the ten configurations after some initial familiarization with the square configuration. Each mouse explored one configuration per day and explored the full sequence of ten configurations every ten days. Each mouse completed two or three full sequences of configurations. From these data (12,606 cell-sequences combined across 7 mice), we computed firing rate maps for each cell-sequence in each configuration by binning and smoothing the firing rate as a function of position. Next, we correlated these rate maps across configurations and across partitions to characterize the representational geometry of mouse CA1 at both resolutions, analogous to our characterization of human spatial memory (Fig. 4b). Of note, the representational geometry of mouse CA1 was consistent across mice and became more similar across experience at both resolutions, indicating that this representational geometry is common across mice and experience at least to the precision of our data.

**Figure 4.**
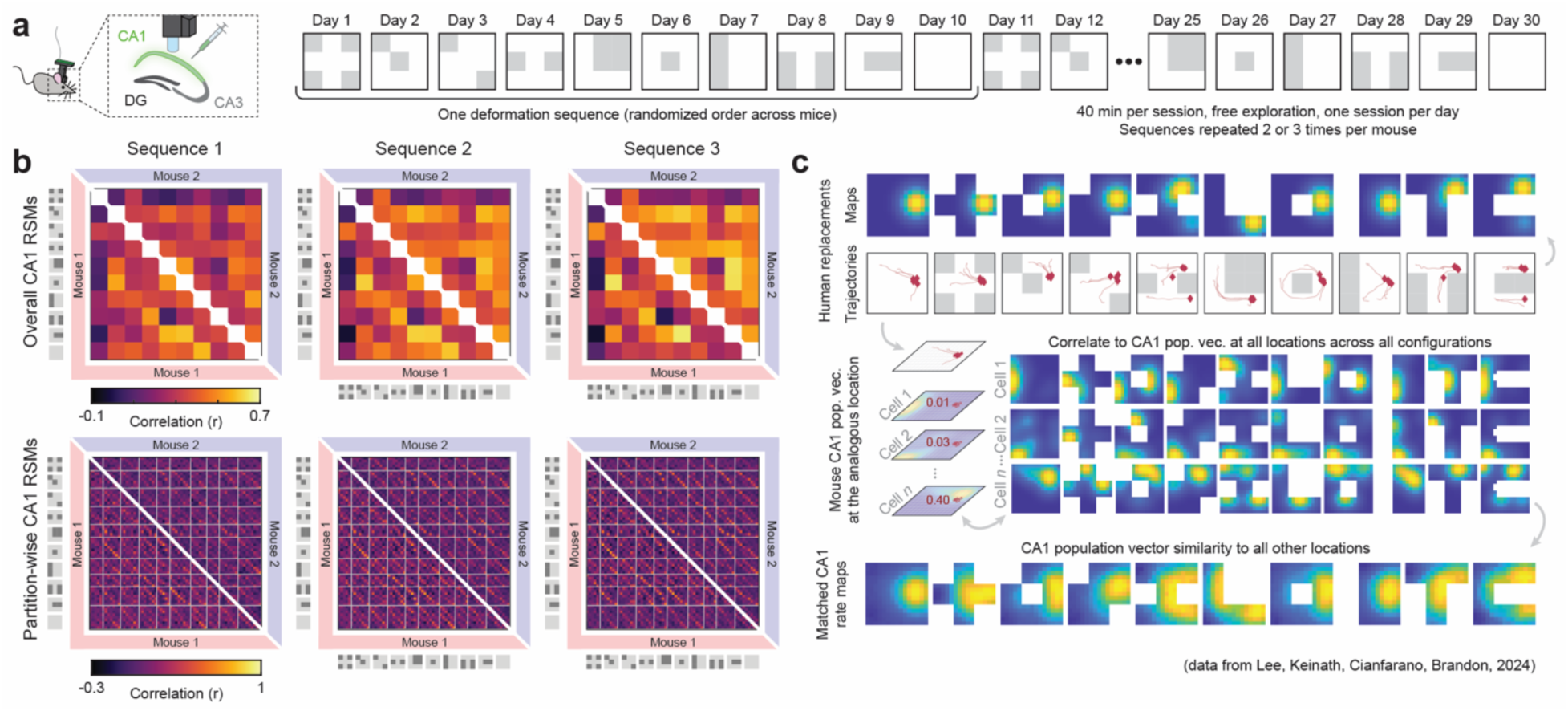
(**a**) Schematic of miniscope imaging of mouse CA1 in the deformation shapespace over weeks of experience. (**b**) Examples of overall RSMs and partition-wise RSMs from two mice across three sequences of the shapespace. Note the consistency across sequences and mice. (c) In order to control for differences in allocation statistics between mouse CA1 and human spatial memory, we computed a matched CA1 rate map for each human replacement map by correlating the CA1 population vector at the analogous pixel location to all other pixels. By varying the CA1 cells which contribute to the population vector, we can test whether the representational geometry of some CA1 subpopulations better resemble that of human spatial memory. RSMs in (**b**) are derived from the raw data, not matched maps.

One key difference between mouse CA1 and human spatial memory assays in these experiments is how replace locations and CA1 firing fields are allocated to the environment. In our human experiments, replace locations are determined by the experimenter to randomly tile the environment, subject to some constraints. In our CA1 data, allocation of firing fields is determined by the hippocampus^54^ with a tendency to overrepresent the boundaries of the environment^55^. As these allocation statistics make a significant contribution to the representational geometry^50^, it is important to equate the two before comparison. To do so, for each replacement map we computed a matched CA1 rate map by comparing the CA1 population vector at the analogous replace location to the CA1 population vectors at all locations across all configurations (see Methods; Fig. 4c). Matched mouse CA1 RSMs were computed from these matched maps. This procedure allows us to control for differences in allocation statistics and also conveniently equates the number of maps used to compute both human and mouse RSMs. Because some cells were silent during some sessions, we also subsampled our data to ensure that all matched maps were derived by correlating equal numbers of active cells for each comparison.

### Mouse CA1 and human spatial memory recapitulate similar representational geometries at the resolution of the entire environment

Do mouse CA1 and human spatial memory recapitulate similar representational geometries in deformed environments? To address this question, we began by computing the rank correlation (Kendall’s *τ*) between the overall human spatial memory RSM (oRSM_HSM_) and the matched mouse RSM (oRSM_CA1_; n = 4,441 active cells per comparison). Prior work demonstrates that this measure is particularly apt when comparing representational geometries^48,56^. Importantly, both RSMs depend to some extent on the degree of map smoothing, and the proper values for these parameters are not known *a priori*. Therefore, we performed a grid search while varying the smoothing of replacement maps and matched CA1 maps (Fig. 5a), and focused further analysis on the maximizing parameterization (*σ_HSM_* = 1 pixel, 0.333 m; *σ*_*CA*1_ = 2.5 pixels, 12.5 cm). This biases us towards observing greater similarity between the two species, and therefore a reliable difference at this maximizing parameterization is strong evidence that the two geometries differ.

**Figure 5.**
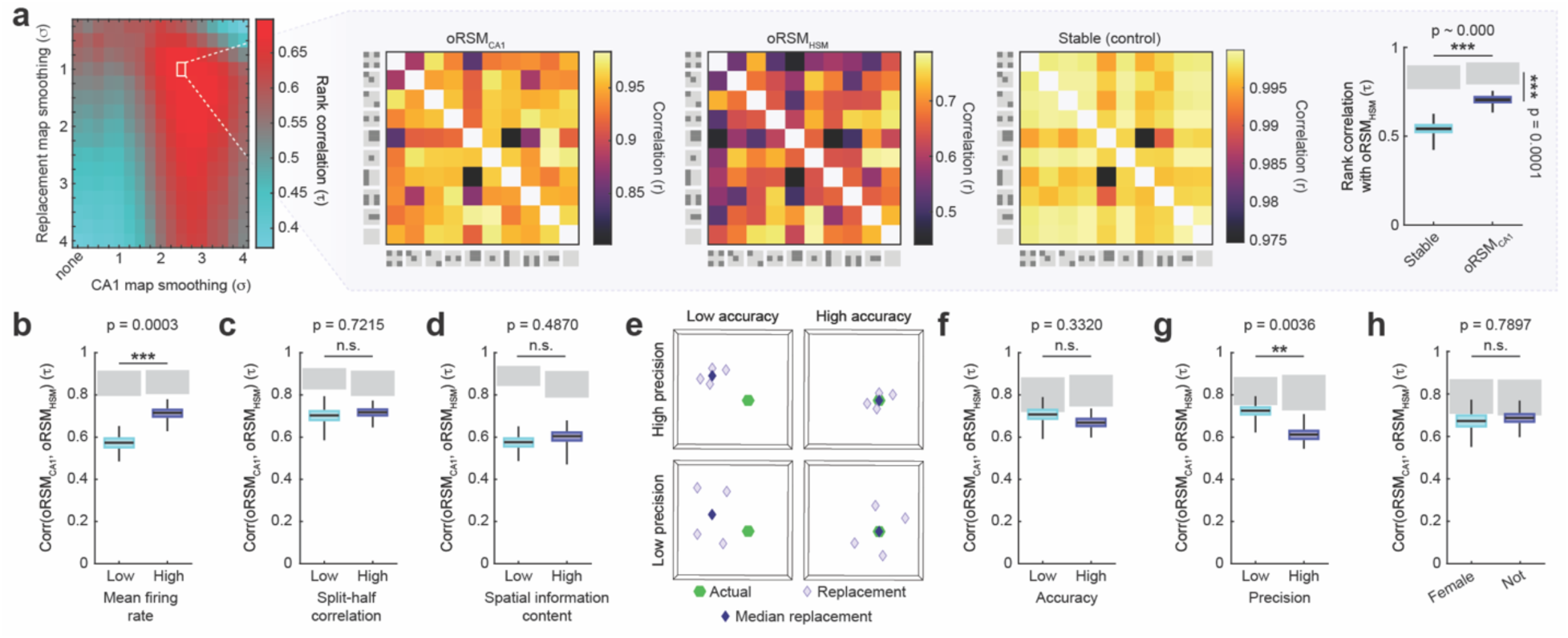
P-values estimated via nonparametric bootstrapping (see Methods). (**a**) At optimal smoothing parameters, matched oRSM_CA1_ and oRSM_HSM_ are highly correlated, exceeding the correlation between oRSM_HSM_ and the stable control. Nevertheless, the true correlation falls significantly below the joint noise ceiling (grey), indicating that oRSM_CA1_ and oRSM_HSM_ do reliably differ. (**b**) Rank correlation between oRSM_HSM_ and oRSM_CA1_ when oRSM_CA1_ is computed only from cells with low or high mean firing rates, determined by median split. (**c**) As in (**b**) except split according to split-half correlations. (**d**) As in (**b**) except split according to spatial information content. (**e**) Schematic of precision and accuracy. (**f**) Rank correlation between oRSM_HSM_ and oRSM_CA1_ when oRSM_HSM_ is computed only from replacements with low or high accuracy, determined by median split. (**g**) As in (**f**) except split according to precision. (**h**) As in (**f**) except split according to the gender of the participants. **p<0.01, ***p<0.001, uncorrected

Both oRSMs were highly correlated (Fig. 5a). For comparison, we computed a stable control RSM from the CA1 data where maps in all configurations were identical to the square except that closed partitions were inaccessible. The rank correlation of the true RSMs significantly exceeded correlation with the minimal variability in this control (Fig. 5a). To better contextualize the high correlation we observed between the oRSM_HSM_ and oRSM_CA1_, we estimated the correlation we should expect if the two were drawn from the same underlying distributions and only decreased due to measurement noise. Were this to be the case, then replacement maps and matched CA1 rate maps should be interchangeable. Thus, we randomly shuffled the assignment of replacement maps and matched CA1 rate maps between the two conditions and recomputed the correlation between the resulting RSMs 500 times. We refer to this distribution as the joint noise ceiling and note that it differs from a traditional noise ceiling^48,56^ (see Methods). If the correlation between oRSM_HSM_ and oRSM_CA1_ was only reduced by measurement noise, then their actual correlation should fall within the joint noise ceiling. If not, then these two RSMs are reliably dissimilar. We found that the actual correlation between oRSM_HSM_ and oRSM_CA1_ fell below the 95% confidence interval of the joint noise ceiling (Fig. 5a), indicating that the two are reliably dissimilar. These results demonstrate that human spatial memory and mouse CA1 instantiate similar, but not identical, representational geometries at the resolution of the whole environment during deformations.

Neural populations in rodent CA1 are known to be heterogenous in their functional properties^52,57–60^. We next asked whether the representational geometry of some subpopulations of cells better resembled that of human spatial memory. To do so, we varied the cells which contributed to the population vector when constructing our match CA1 rate maps (Fig. 4c). Note that this allows us to vary which cells contribute to the resulting matched CA1 representational geometry without reducing the number of matched maps. Dividing by median split and controlling for the number of active cells contributing to each comparison, we found that the representational geometry of cells with higher mean firing rates better resembled that of human spatial memory (Fig. 5b; n = 1088 active cells per comparison). Cells with higher split-half correlations – a measure of spatial tuning stability – and high spatial information content – a measure of spatial tuning specificity – also exhibited numerically better resemblance with human spatial memory, although these differences did not reach significance (Fig. 5c,d; split-half correlations, n = 1149 active cells per comparison; spatial information content, n = 943 active cells per comparison).

Humans also exhibit individual differences in their spatial memories and navigational strategies^61,62^. Thus, we next asked whether the representational geometry of some replacements better resembles that of mouse CA1. We considered two factors which might vary across replacements: accuracy and precision (Fig. 5e). Accuracy was quantified as the inverse distance between the median replacement location and the actual location of the object in the square configuration during the deformation blocks. Precision was quantified as the inverse replacement spread in the square configuration during the deformation blocks. The rank correlation between oRSM_HSM_ and oRSM_CA1_ did not significantly depend on replacement accuracy (Fig. 5f). However, the rank correlation between oRSM_HSM_ and oRSM_CA1_ did significantly depend on replacement precision. Low-precision replacements recapitulated a representational geometry which better resembled that of mouse CA1 relative to their high-precision counterparts (Fig. 5g). Finally, the rank correlation between oRSM_HSM_ and oRSM_CA1_ did not significantly on the gender of our participants.

### The geometry of human spatial memory resembles a change-resistant version of that of mouse CA1 at the resolution of individual partitions

In the previous section we compared the representational geometries of mouse CA1 and human spatial memory in deformed environments at the resolution of the entire map. Our partition-based parametric design also allows us to compare representational geometry at a finer resolution. We began by computing the rank correlation between the partition-wise human spatial memory RSM (pRSM_HSM_) and the matched mouse RSM (pRSM_CA1_). To determine which smoothing parameters maximized the similarity of these RSMs, we again performed a grid search while varying the smoothing of replacement maps and matched CA1 maps (Fig. 6a). At the maximizing parameterization (*σ_HSM_* = 2.25 pixels, 0.75 m; *σ*_*CA*1_ = 1.75 pixels, 8.75 cm), these RSMs were highly correlated (Fig. 6a). For comparison, we considered a control whereby each partition was identical only to itself across all configurations, as might be expected if the deformations had no impact on the spatial code within accessible partitions. The rank correlation of the true RSMs significantly exceeded correlation with this control (Fig. 6a). Nevertheless, the rank correlation of the true RSMs fell significantly short of the joint noise ceiling, indicating a reliable difference between these two representational structures.

**Figure 6.**
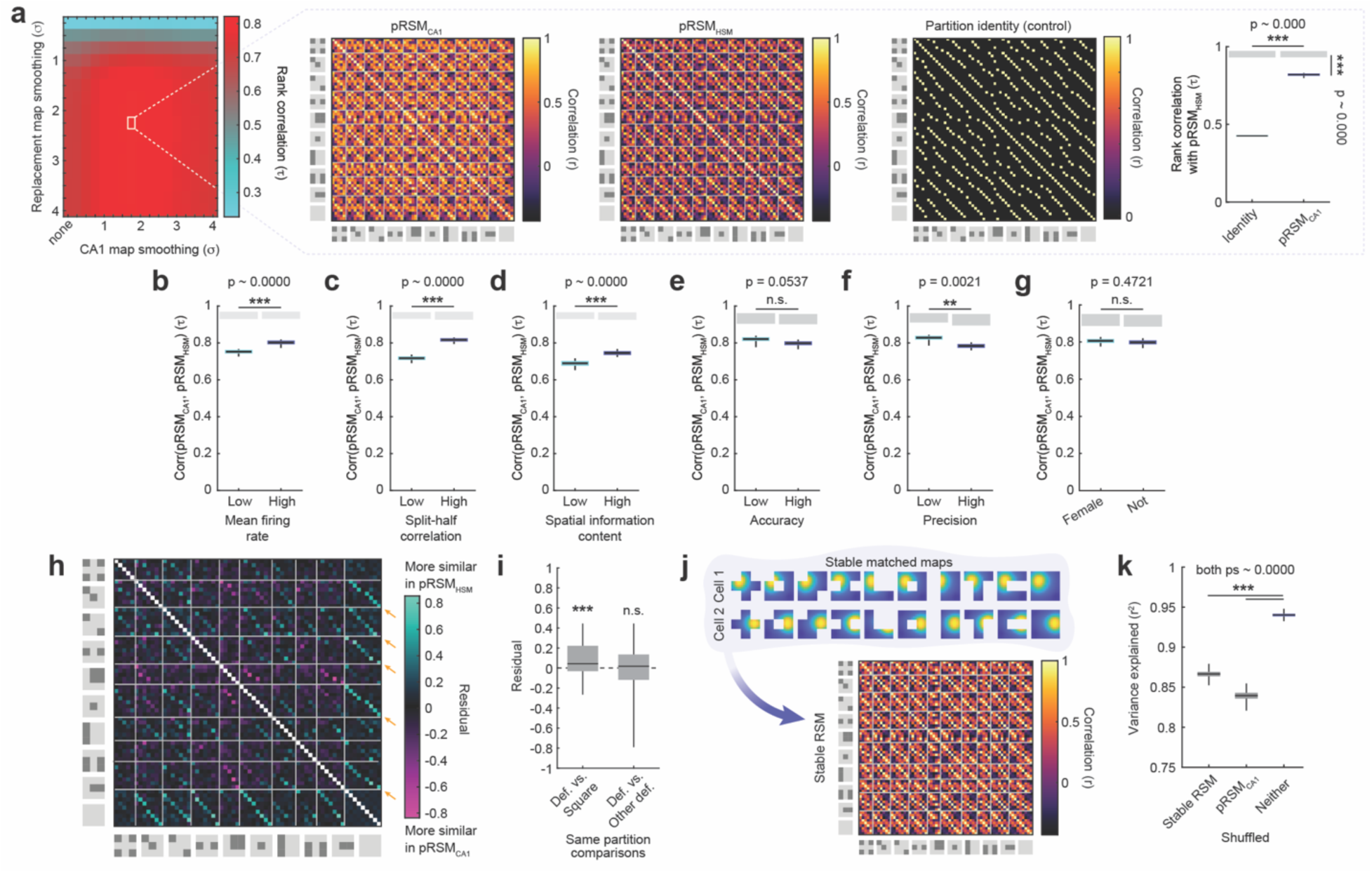
P-values estimated via nonparametric bootstrapping unless otherwise noted (see Methods). (**a**) At optimal smoothing parameters, matched pRSM_CA1_ and pRSM_HSM_ are highly correlated, exceeding the correlation between pRSM_HSM_ and the partition identity control. Nevertheless, the true correlation falls significantly below the joint noise ceiling (grey), indicating that pRSM_CA1_ and pRSM_HSM_ do reliably differ. (**b**) Rank correlation between pRSM_HSM_ and pRSM_CA1_ when pRSM_CA1_ is computed only from cells with low or high mean firing rates, determined by median split. (**c**) As in (**b**) except split according to split-half correlations. (**d**) As in (**b**) except split according to spatial information content. (**e**) Rank correlation between pRSM_HSM_ and pRSM_CA1_ when pRSM_HSM_ is computed only from replacements with low or high accuracy, determined by median split. (**f**) As in (**e**) except split according to precision. (**g**) As in (**e**) except split according to the gender of the participants. (**h**) Residual when explaining the variance in pRSM_HSM_ with pRSM_CA1_ as a predictor. Note the bands of teal for comparisons of the same partition between deformed configurations and the square configuration (orange arrows). (**i**) Residual for comparisons of the same partition between deformed configurations and the square configuration, or deformed configurations and other deformed configurations. Significance markers indicate outcome of signed-rank tests versus 0. (**j**) A stable RSM was derived from stable matched CA1 maps, where all configurations are identical to the square except that some partitions are inaccessible. (**k**) Variance in pRSM_HSM_ explained by a combination of pRSM_CA1_ and the stable RSM when both are intact versus when one predictor is shuffled. **p<0.01, ***p<0.001, uncorrected

We next asked whether the representational geometry of certain subpopulations of CA1 cells better resembled that of human spatial memory at this finer resolution. Dividing by median split, we found that the representational geometry of cells with higher mean firing rates (n = 1088 cells per comparison), split half correlations (n = 1108 cells per comparison), and spatial information content (n = 1074 cells per comparison) all better resembled that of human spatial memory (Fig. 6b-d). These results match our findings when comparing overall RSMs, and collectively demonstrate that the CA1 subpopulations with geometries which best resemble that of human spatial memory are consistent across scales of analysis.

We then asked whether the rank correlation between pRSM_HSM_ and pRSM_CA1_ depended upon replacement accuracy or replacement precision. Similar to our findings at the resolution of the entire map, we found that the cross-species resemblance in representational geometry did not significantly depend on the accuracy of replace locations (Fig. 6e), but did significantly depend on precision (Fig. 6f). Again, low precision replacements yielded geometry which better resembled that of CA1. Finally, the rank correlation between pRSM_HSM_ and pRSM_CA1_ did not significantly depend on the gender of our participants. These results thus demonstrate that the geometry of replacements which best resemble that of mouse CA1 are consistent across resolutions of analysis.

Despite the high rank correlation between pRSM_HSM_ and pRSM_CA1_, comparison with the joint noise ceiling indicates that the two reliably differ. To gain insight into the difference between these two representational structures, we turned to a general linear model (GLM) approach. We first Fisher-transformed both RSMs to overcome the challenge of extreme skew at high correlation values. Next, we utilized a GLM to explain the variance in pRSM_HSM_ with pRSM_CA1_ as a predictor. Finally, we examined the residuals of this prediction to determine where these RSMs diverged (Fig. 6h). We observed that comparisons between the same partition in deformed versus square configurations were more similar in the pRSM_HSM_ than expected from the pRSM_CA1_ (Fig. 6h, teal bands highlighted by orange arrows). Quantifying this, we found that comparing the same partition between deformed and square configurations yielded significantly positive residuals (signed-rank test versus 0: Z(47943) = 0.2851, p = 0.7755), whereas comparisons of the same partitions among deformed configurations did not significantly deviated from zero (signed-rank test versus 0: Z(47943) = 0.2851, p = 0.7755; Fig. 6i). This suggests that human spatial memory is more resistant to the representational changes induced by environmental deformations than mouse CA1.

To more directly test this possibility, we repeated our GLM analysis to explain the variance in the pRSM_HSM_ except including two predictors: pRSM_CA1_ and an additional stable RSM derived from the CA1 data where maps in all configurations were identical to the square except that closed partitions were inaccessible (Fig. 6j). We found that combining these two predictors explained a substantial portion of the variance in the pRSM_HSM_ (∼94%) and that the two together explained significantly more variance than when either of these predictors were shuffled (Fig. 6k). Together, these results indicate that the representational geometry of human spatial memory during environmental deformations closely resembles a change-resistant version of mouse CA1.

### When humans navigate a larger space, local impact scales up and representational geometry is preserved

For both human spatial memory and rodent hippocampal representations, the scale of the environment is an important determinant of many representational properties^54,55,63^. Thus it is possible that the cross-species similarity we observed in our first experiment is particular to the scale of the environments explored by both navigators. On the other hand, it is possible that the impact of environmental deformations on representational structure is invariant to the scale of the environment. While a full contrast of environmental scales in both rodents and humans is beyond the scope of this work, we began to address these possibilities by repeating our human spatial memory experiment in a new cohort of participants (n = 102) in a 10 x 10 m virtual environment, double the length and width of the environment in our first experiment (Fig. 7a). All other design elements were identical. Participants moved more quickly but were less accurate during the deformation blocks in the larger environment (Fig. 7b,c).

**Figure 7.**
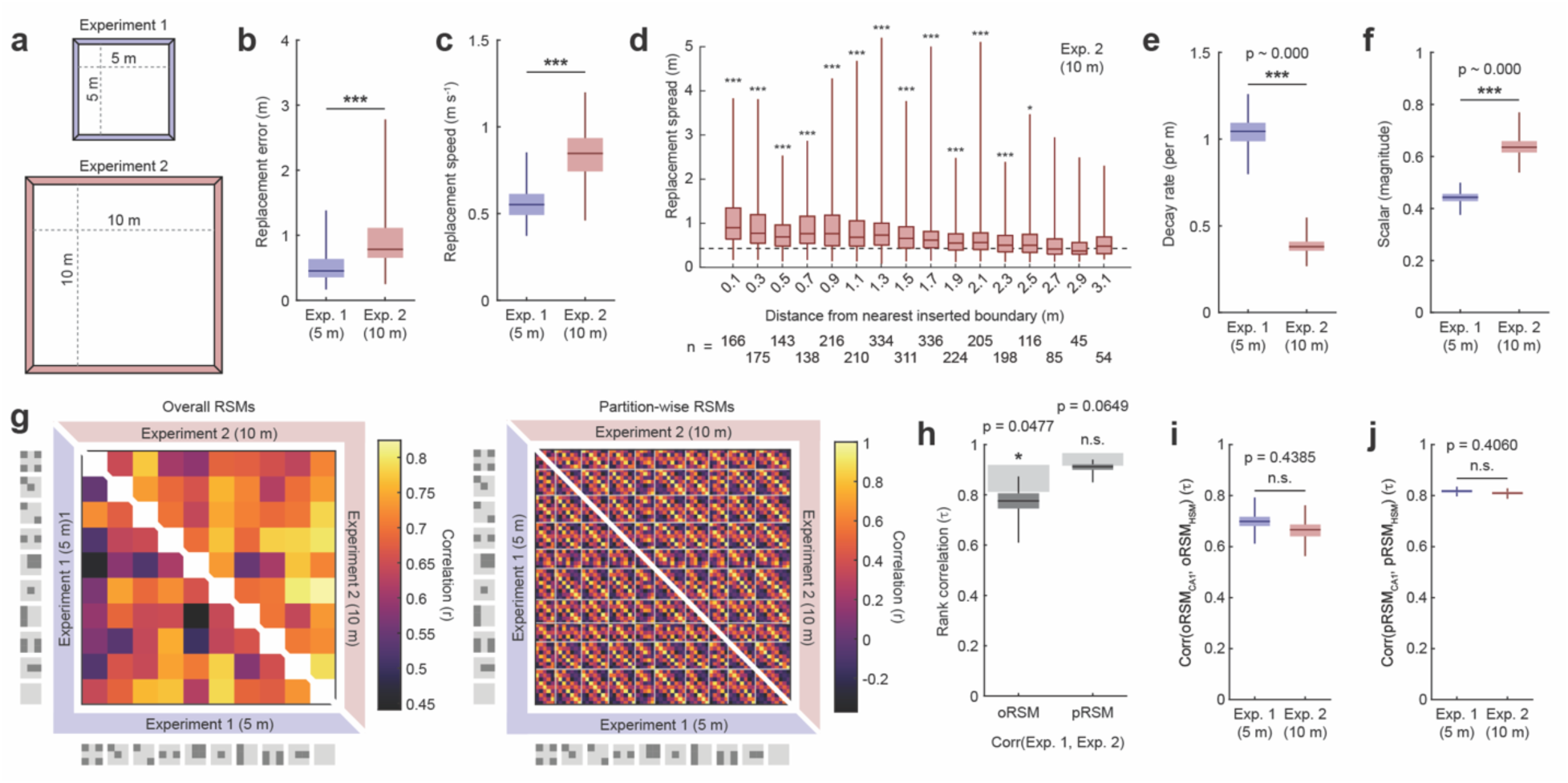
(**a**) Schematic of the difference in environment size between experiments one and two. (**b**) Replacement error was larger in the larger environment (rank-sum test: Z(7366) = -7.9423, 1.9841e-15). (**c**) Replacement speed was faster in the larger environment (rank-sum test: Z(6113) = -10.8715, 1.5764e-27). (**d**) Replacement spread as a function of distance to the nearest inserted boundary in the larger environment, aggregated across configurations for accessible objects. Significance markers indicate the outcome of a rank-sum test between the replacement spread of the denoted distance group and the square configuration. (**e**) Rate of decay of replacement spread as a function of distance from the nearest inserted boundary, determine by fitting a two-parameter exponential decay function (see Methods). (**f**) As in (**e**) except for the magnitude of replacement spread. (**g**) Comparison of oRSMs and pRSMs between the smaller and larger environments (**h**) Rank correlation between the RSMs of the smaller and larger environments. P-values estimated via nonparametric bootstrapping (see Methods). (**i**) Rank correlation between the oRSM_HSM_ and oRSM_CA1_ for the smaller and larger environments. P-values estimated via nonparametric bootstrapping (see Methods). (**j**) As in (**i**) except between the pRSM_HSM_ and pRSM_CA1_ for the smaller and larger environments. *p<0.05, ***p<0.001

We first asked whether the local impact of deformations on replacement spread differed as a function of environment scale. As before, for each object we computed the replacement spread observed in each configuration, as well as the distance of the median replacement location in the square to the nearest insert boundary. We then aggregated accessible objects across deformed configurations. In the larger environment, we again observed a local impact of deformations on replacement spread, with objects nearer to inserted boundaries exhibiting larger spread. Notably, in this larger environment we observed significantly elevated replacement spread relative to that of the square configuration for objects replaced within 2.5 m of the nearest boundary (Fig. 7d) – double the distance we observed in the 5 x 5 m environment. To quantify this difference more precisely, we fit a two-parameter exponential decay function to the replacement spread based on the distance to the nearest inserted boundary for both experiments (Fig. S1; see *Methods*). The two parameters in this function allowed us to flexibly capture differences in both the magnitude of replacement spread change and rate of replacement spread decay with increasing distance. This analysis revealed that the impact of deformations in the large environment perseverated at distances farther from the inserted boundaries, as reflected by a decay rate roughly half that of the smaller environment (Fig. 7e). Moreover, deformations in the larger environment had a greater impact on replacement spread in general, as indicated by a greater scalar parameter (Fig. 7f). Together, these findings demonstrate that the local impact of environmental deformations does depend on the scale of the environment.

It is possible that the differences we observe in the local impact of deformations between environments of different scale yield differences in representational geometry. On the other hand, it is possible that these differences act to preserve the representational geometry across these scales. To address these possibilities, we first computed replacement maps for the data from our larger environment, doubling our bin size so that replacement maps were equal in pixel size for both the smaller and larger environments (15 pixels x 15 pixels). Next, we computed overall and partition-wise RSMs for this larger environment and compared these with the RSMs from the smaller environment (Fig. 7g). Both oRSMs and pRSMs exhibited rank correlations approaching the joint noise ceiling (Fig. 7h), indicating that the representational geometry is as similar across these two scales as would be expected if the two were drawn from identical distributions and only impacted by measurement noise. Moreover, both the smaller and the larger environment RSMs were equally correlated with those of mouse CA1 (Fig. 7i,j). This indicates that the cross-species resemblance is tolerant to differences in the scale of the environment that human participants navigated. Altogether, these results demonstrate that the local impact of deformations on human spatial memory scales with the size of the environment, preserving the representational geometry and cross-species resemblance, at least on the range tested here.

### Representational geometry is preserved when distal visual input is masked during retrieval

Our previous findings indicate that the impact of environmental deformations on human spatial memory resembles a change-resistant version of mouse CA1. While there could be many reasons for this cross-species difference in representational structure, one possibility is that the human visual advantage plays a key role^44,64^. If so, then reducing the human visual advantage during deformation trials might lead the representational geometry of human spatial memory to better resemble that of mouse CA1. To address this possibility, we carried out a third VR experiment in a 5 x 5 m virtual environment where visual fog reduced the human visual advantage during replacement trials (n = 105 participants; Fig. 8a). Most design elements were identical to those of experiment one except that an additional block of replacement and collection trials was carried out to introduce participants to the experience of replacing objects with fog prior deformation trials (Fig. 8a). Participants accuracy and replacement speeds were impacted by the introduction of fog but recovered during deformation blocks (Fig. 8b,c).

**Figure 8.**
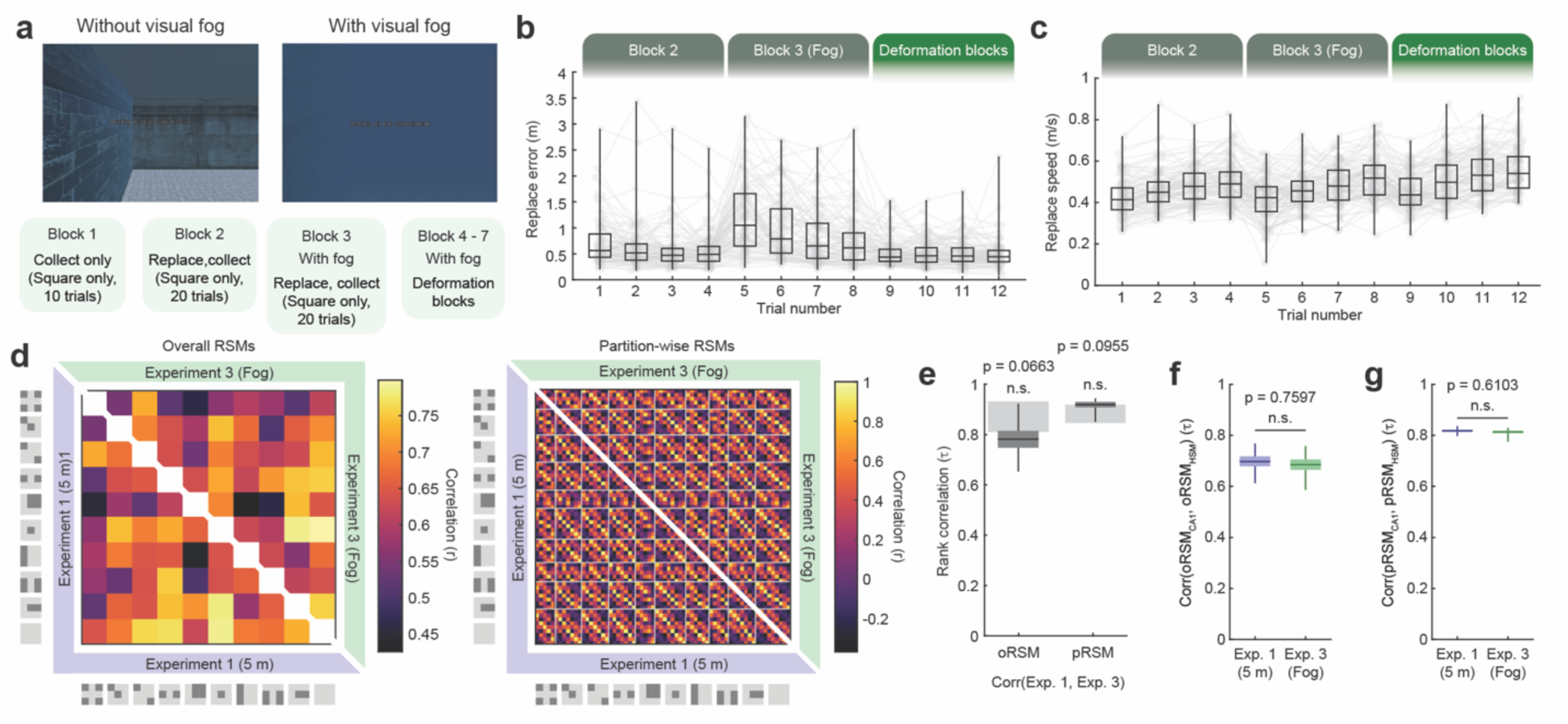
P-values estimated via nonparametric bootstrapping (see Methods). (**a**) Schematic of block design for experiment three, with screenshots with and without visual fog. (**b**) Error in replace locations in the square configuration across blocks. (**c**) Replacement speed in the square configuration across blocks. (**d**) Comparison of oRSMs and pRSMs between experiments one and three, without and with visual fog during replacements. (**e**) Rank correlation between the RSMs of experiments one and three, without and with visual fog during replacements. (**g**) Rank correlation between the oRSM_HSM_ and oRSM_CA1_ for experiments one and three, without and with visual fog during replacements. (**g**) Rank correlation between the pRSM_HSM_ and pRSM_CA1_ for experiments one and three, without and with visual fog during replacements.

We first asked whether reducing the human visual advantage during retrieval impacted the representational geometry of human spatial memory. We compared overall and partition-wise RSMs between this experiment and those of experiment one, where the environment was of comparable size but visual access remained unaltered during replacement trials (Fig. 8d). Rank correlations of both oRSMs and pRSMs between the two experiments approached the joint noise ceiling, indicating that the representational geometry was as similar as would be expected if the two were drawn from identical distributions and only impacted by measurement noise. Next, we asked whether cross-species resemblance was impacted by the reducing the human visual advantage. Comparing both sets of RSMs with those of mouse CA1 revealed that human spatial memory equally resembled mouse CA1 with and without the introduction of visual fog. Together, these results demonstrate that neither the representational geometry of human spatial memory as assayed here nor its resemblance with mouse CA1 depends on the human visual advantage during retrieval.

## Discussion

Extensive prior work has established foundational qualitative similarities between the ways in which rodent neural representations and human spatial memory are impacted by environmental manipulations. Here, we exploited recent technical and theoretical advances to quantitatively characterize the representational structure of human spatial memory during environmental deformations and compare it with that of mouse CA1. Leveraging immersive VR in humans, we find that deformations induce local distortions to human spatial memory which mimic those observed in mouse CA1. These distortions yield a representational geometry which closely resembles a change-resistant version of that of mouse CA1. The representational geometry of mouse CA1 subpopulations with higher firing rates, spatial tuning stability, and spatial specificity all better resembled the geometry of human spatial memory. The precision, but not accuracy, of human spatial memory also influenced cross-species resemblance, with the geometry of low-precision memories better resembling that of mouse CA1. Comparing humans navigating smaller and larger versions of the same deformed environments, we found that the local impact of deformations scaled with the size of the environment, which in turn preserved the representational geometry and cross-species resemblance across environmental scales. Finally, the geometry of human spatial memory was not impacted by reducing visual acuity during retrieval, suggesting that the degree of cross-species resemblance was not dependent upon the human visual advantage. These results go beyond traditional qualitative comparisons to establish common cross-species resemblance in the representational structures of mouse CA1 and human spatial memory during environmental deformations, with a notable difference in the resistance to change between these assays.

The representational geometries of human spatial memory and mouse CA1 primarily differed in their resistance to change during deformations. There may be many reasons for this difference, even assuming identical hippocampal representations in the two species. One possibility is that CA1 subpopulations differentially inform memory-guided behaviors. For example, memory-guided behaviors might rely more heavily on cells with higher firing rates, spatial tuning specificity, and/or spatial tuning stability, each of which yielded geometries which better resembled that of human spatial memory. It is also possible that the memory-guided behavior we assay here in humans is the product of multiple memory systems and/or multiple navigational strategies^61–63^, some of which differ in their representational geometry. For example, contributions from prefrontal cortex or the striatum might complement or compete with that of the hippocampus^51,65,66^. Decision-making processes downstream of the hippocampus may also play an important change-resisting role^67,68^. Of course, it is alternatively possible that the differences we observe here reflect nuanced differences in hippocampal representational geometry, consistent with our growing appreciation for cross-species differences in these neural codes.

The datasets we compare here were specifically collected to enable comparison of representational geometry across techniques and species^47^. One advantage of such designs is that the data can continue to be leveraged in future comparisons. For example, future work might leverage this design to characterize changes across the lifespan in the representational geometry of aging humans, in both typical aging and diseases of memory such as Alzheimer’s^69–71^. Computational work might address whether common models can explain representational geometry in both mouse CA1 and human spatial memory^72^, perhaps focusing specifically on the structure shared between the species. Characterizing the representational geometry of other navigationally-relevant brain regions such as entorhinal, prefrontal, retrosplenial, and posterior parietal cortices as well as the striatum in rodents might address whether contributions from these other structures could explain change-resistance at the level of behavior. Finally, assaying spatial memory in mice might address whether change-resistance at the level of behavior is common across species or reflects a fundamental difference in the representational structure of cognitive maps. Together, this approach can help us leverage diverse assays to tease apart nuanced alternatives, and thus better understand representational differences across brain regions, species, levels of explanation, and health statuses.

## Methods

### Human participants

Experiment 1 included n = 104 participants, with a gender distribution of female = 62, male = 42, and non-binary = 0 and an age distribution of = 20.308 ± 3.268 years (mean ± standard deviation) ranged 18 to 36 years. Four additional participants were not included in the dataset because they did not outperformed chance during familiar square replacements or walked through virtual walls. Experiment 2 included n = 105 participants, with a gender distribution of female = 77, male = 27, and non-binary = 1 and an age distribution of = 19.673 ± 2.338 years (mean ± standard deviation) ranged 18 to 30 years. Three additional participants were not included in the dataset because they did not outperformed chance during familiar square replacements or walked through virtual walls. Experiment 3 included n = 102 participants, with a gender distribution of female = 66, male = 36, and non-binary = 0 and an age distribution of = 20.188 ± 3.405 years (mean ± standard deviation) ranged 18 to 34 years. Four additional participants were not included in the dataset because they did not outperformed chance during familiar square replacements or walked through virtual walls. All participants provided informed consent in accordance with the Institutional Review Board of the University of Illinois Chicago and were paid or received course credit for their participation in these experiments. Sample size was chosen prior to conducting this experiment such that the total number of replaced objects aggregated across participants was approximately equal to the number of hippocampal cells typically assayed within a mouse in the mouse CA1 dataset^50^.

### Human data acquisition

All experiments were carried out on an HTC VIVE XR Elite head-mounted display with a resolution of 1920 x 1920 pixels per eye yielding a field of view of approximately 110° and a refresh rate of 90 Hz. Prior to the experiment, interpupillary distance and focus for both eyes were adjusted for each participant. Experiment 1 was run by streaming the developed application over dedicated WiFi on an ASUS ROG Rapture GT-AX11000 router using SteamVR (v2.9.6) and the VIVE streaming hub (v2.3.3b). Experiments 2 and 3 were run via streaming in some participants (Experiment 2: 72 participants; Experiment 3: 44 participants) and via local execution on the head-mounted display in the remaining participants. Streaming led to occasional stutter (approximately once per session) where the headset display went dark for ∼2 seconds before the experiment was restored. Local execution on the head-mounted display greatly reduced these events (less than once every ten participants). Experiment 1 took 48.4 ± 9.6 min (mean ± standard deviation) to complete, Experiment 2 took 58.5 ± 7.9 min (mean ± standard deviation) to complete, and Experiment 3 took 61.8 ± 10.0 min (mean ± standard deviation) to complete. Experiments were typically completed on a single battery charge, but if batteries needed to be replaced this happened between blocks. Throughout the experiment, the position and heading in all three dimensions relative to the virtual environment were recorded at a rate of 90 Hz in a comma separated file written either to the streaming desktop or to the local storage of the headset. Participant responses were registered by pressing the primary A button on the VIVE remote or an analogous handheld remote. All analyses were derived from these data.

### Human virtual environments

All experiments were developed in the Unity game engine (2022.3.8f1; Unity Technologies) by the experimenters. In all experiments, the virtual environment was comprised of four walls with repeating textures, one of which was textured differently to provide an orientational cue. The ground was textured with repeating tile. Textures were repeating to minimize their utility as location cues. All walls were 2 m in height and the environment was surrounded by a skybox in which the sun and clouds provided orientational cues rendered at infinity. In Experiment 3, the environment was encased in a grey-textured box because the visual fog mechanic did not obscure the skybox during testing.

During deformation blocks, the environment was deformed on certain trials by walls which rose up from below the environment. All walls were textured to match the walls of the environment and were connected by a roof which matched the texture of the floor. Walls took approximately 3 s to rise into place, calibrated to be as prompt as possible without startling the participants. When deformation trials in that configuration were complete, the walls retracted into the floor within approximately 1 s. Participants were instructed to treat the virtual walls as if they were physical walls, and any participant who chose to violate this by walking through a wall was excluded from the experiment.

Participants learned the locations of five objects: a box, barrel, axe, broom, and saw. Each object was derived from a freely available asset in the Unity Asset Store. At times, participants were also guided by a waypoint, which was a metallic purple floating capsule. When visible in the environment, objects and the waypoint floated centered around 1.25m above the ground and slowly rotated at ∼9°/s. Objects were randomly distributed throughout the environment subject to the constraints that all objects must be at least a minimum distance from the outer walls of the environment (to protect participants from accidentally contacting the real-world bounds of the environment) and a minimum distance from one another. Waypoint locations were randomly distributed subject to the constraint that they were at least the minimum distance away from all walls and would not lead the participant to a to-be-enclosed location on the next trial during deformation blocks.

In the third experiment, visual fog occluded the distal visual world during some trials. This fog saturated exponentially and fully occluded the visual world beyond 0.5 m from the participant. Fog appeared and disappeared gradually within about 1 s at the beginning and end of those trials.

All instructions (i.e. “Replace the barrel”, “Go to the waypoint”, “Collect the box”) appeared in white text centered in the field of view at a distance of approximately 0.75 m from the participant. These instructions were not obscured by the visual fog.

### Human experimental design

All experiments followed a blocked design modeled on prior work^25,31,51^. In the first block, participants learned the locations of each object by collecting each object twice in a random order while the environment was in the square configuration. In the second block, participants replaced each object four times in the square configuration. Participants were instructed to stand where they felt the object should be and press the button on their handheld remote. After each replacement, participants were given the opportunity to collect that object again to refresh their memory. After each replacement trial, participants were instructed to navigate to a waypoint to encourage them to continue to learn the locations of the objects from different starting points. In Experiment three, participants completed an additional block of square replace and collect trials that was identical to their previous block except that visual fog occluded the distal visual world on replace trials. This block was included to allow participants to gain experience with the fog manipulation prior to the deformation blocks. Finally, in all experiments the last four blocks consisted of deformation trials with a similar structure. In each deformation block, the participants replaced each object in each environmental configuration once (nine deformed configurations and the familiar square configuration). The order of configurations within a block was random, and all five objects were replaced one after another during each configuration to minimize the number of times that the environment changed shape while the participant navigated. Between configuration changes, the participant was guided by a waypoint to a random location that was sampleable in both the current and upcoming configuration. Participants also received three additional chances to collect the objects in the square configuration to refresh their memories, once at the beginning of the block, once after the third configuration, and once after the sixth configuration. Between all blocks the experimenter checked in with the participant to make sure that they were not motion-sick and to provide instructions for the upcoming block. Participants were permitted to take breaks between blocks if they chose.

### Mouse data acquisition

Although the mouse CA1 data analyzed here were previously reported^50^ and are publicly available at [https://zenodo.org/records/14867736], for completeness we describe key details of their acquisition here. Right hippocampal CA1 of naïve mice (C57Bl/6, Charles River; 3 female) was transfected with the calcium indicator GCaMP6f via the viral construct AAV9.syn.GCaMP6f.WPRE.SV40 (Addgene) diluted in sterile PBS. Following transfection, mice were implanted with a 1.8mm diameter GRIN lens (Edmund Optics) above right CA1 (Referenced to bregma: ML = 2.0 mm, AP = -2.1 mm; Referenced to brain surface: DV = -1.35 mm). Implantation required aspiration of intervening cortical tissue. Following implantation, an aluminum baseplate was cemented to the implant to secure the miniscope for imaging.

In vivo calcium videos were recorded with a UCLA miniscope^73^ (v3; miniscope.org) containing a monochrome CMOS imaging sensor (MT9V032C12STM, ON Semiconductor) connected to a custom data acquisition (DAQ) box (miniscope.org) with a lightweight, flexible coaxial cable. The DAQ was connected to a PC with a USB 3.0 SuperSpeed cable and controlled with Miniscope custom acquisition software (miniscope.org). The outgoing excitation LED was set to between 2-8% (∼0.05-0.2 mW), depending on the mouse to maximize signal quality with the minimum possible excitation light to mitigate the risk of photobleaching. Gain was adjusted to match the dynamic range of the recorded video to the fluctuations of the calcium signal for each recording to avoid saturation. Behavioral video data were recorded by a webcam mounted above the environment. The DAQ simultaneously acquired behavioral and cellular imaging streams at 30 Hz as uncompressed avi files and all recorded frames were timestamped for post-hoc alignment.

Calcium imaging data were preprocessed prior to analyses via a pipeline of open source MATLAB (MathWorks; version R2015a) functions to correct for motion artifacts^74^, segment putative cells, and extract transients^75,76^. Autoregressive deconvolution of order two was then used to infer the spiking activity which gave rise to each trace, yielding a nonnegative vector which scaled with the likelihood of spiking during each frame^77^. This vector was treated as the firing rate for each cell. Cells were tracked across sessions on the basis of prominent landmarks, their spatial footprints, and/or centroids^78^. Position of the head was estimated from the behavioral video via DeepLabCut^79^ and interpolated to match the timestamps of the corresponding imaging frames. All further analysis was conducted on the firing rate and position vectors.

Mice freely explored one environment per day for 40 min. The full square environment was 75 cm x 75 cm, surrounded by 60 cm tall external walls and with a blue Lego baseplate affixed at the top-center of the eastern wall to serve as an orienting visual feature. During recording, the environment was dimly lit. All mice explored the square configuration for four sessions prior to the first deformation sequence for familiarization; these initial sessions were not included in our analysis. The order of the environmental configurations was randomized for each mouse, but repeated across sequences such that repeated configurations were always equidistant in time. Each mouse completed two or three sequences. Data were combined across sequences. Because most cells were active during at least some sessions across sequences, most cells likely contributed multiple cell-sequences to the analysis here.

### Quantifying local distortions in human spatial memory

For all analyses, the position and heading data recorded at a rate of 90 Hz were loaded into MATLAB (R2024b) and analyzed using custom written scripts. For each replacement trial, the replacement location was taken as the final location of the headset when the participant pressed the button on the handheld remote.

During deformation blocks, replacement spread was computed as the mean distance of all four object replacements to the median replace location in the square configuration. Median was chosen to reduce the impact of outliers, though similar results are observed with other measures of central tendency such as the mean. Because the median square replace location of some objects was inaccessible in some deformed configurations, only accessible objects were included in the analysis.

To characterize the change in replacement spread as a function of distance to the nearest inserted boundary, in some cases we fit a two-parameter exponential decay function to these data. To do so, we first subtracted the median replacement spread in the square configuration, so that this function should decay to zero. Next, aggregating our accessible replacement data across all deformed configurations and participants, we fit the function:

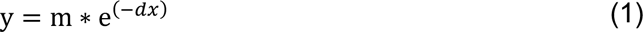

where *x* is the distance to the nearest inserted boundary, *d* is the rate of exponential decay (Fig. 7e), m is a scalar which captures differences in the magnitude of this function (Fig. 7f), and y is the resulting spread. To estimate the variability in parameter fits, we recomputed fits over 500 resamplings with replacement of the data prior to function fitting. Significance markers and p-vales were estimated via nonparametric comparison across these 500 bootstrapped samples.

### Human replacement maps

Replacement maps were computed for each object and configuration by dividing the environment into a 15 pixel x 15 pixel grid. The value of each pixel was then set to the proportion of times the object was placed at that location across all four deformation blocks. These maps were then smoothed with an isotropic Gaussian filter with standard deviation corresponding to the parameterizations in the main text. Locations that were inaccessible in a given configuration were excluded from further comparison.

### Mouse CA1 rate maps

Rate maps were computed for each cell-sequence and configuration by dividing the environment into a 15 pixel x 15 pixel grid. The value of each pixel was then set to the mean firing rate of that cell while the mouse was at that pixel location. These maps were then smoothed with an isotropic Gaussian filter with standard deviation corresponding to the parameterizations in the main text. Locations that were inaccessible in a given configuration were excluded from further comparison.

### Matching mouse CA1 rate maps to human replacement maps

In order to control for differences in allocation statistics between mouse CA1 and human spatial memory, we computed a matched CA1 rate map for each human replacement map as follows. First, for a given object and all CA1 cells we wished to include, we computed the dot product between the unsmoothed square replacement map for that object and each cell’s smoothed square CA1 rate map. This yielded the CA1 population vector expected to correspond to that object’s replacement map. Next, we correlated this population vector across all pixels in all configurations, yielding a matched CA1 map for each configuration for that object. By varying the CA1 cells which contribute to this procedure, we can test whether the representational geometry of some CA1 subpopulations better resembles that of human spatial memory, while matching the number of maps and allocation statistics of those maps. A graphical representation of this procedure can be found in Figure 4c.

### Comparing representational geometries

Previous work has demonstrated through simulation that rank correlation (Kendall’s Tau) is the preferred metric when comparing RSMs, as this measure converges on the true model even at low sample sizes; therefore we use this metric when comparing RSMs in most cases.^48,56^ We note that similar results are observed with Pearson correlations as well.

To bound the correlations when comparing two RSMSs, we also computed a joint noise ceiling. This joint noise ceiling is a nonparametric estimate of the rank correlation we should expect to observe between the mean RSMs of both groups if the two groups were drawn from an identical distribution and only affected by measurement noise. If this were the case, then group identity should be arbitrary. Thus, to estimate the joint noise ceiling, we shuffled group identity and recomputed the rank correlation between the mean RSMs of both groups 500 times. Actual rank correlations were nonparametrically tested against this distribution to determine whether or not the groups significantly differed.

We note that the joint noise ceiling we compute here differs from the traditional noise ceiling of Representational Similarity Analysis^48,56^. The traditional noise ceiling takes into account the variability of a single group and bounds the correlation that one should expect to observe between a new individual and the existing group average. In the human VR experiments reported here, each participant replaced only five objects, thus any RSM computed at the level of the participant would be highly variable, failing to capture the reliable representational structure at the level of the population. As such, the traditional noise ceiling is dispreferred here. Were we to have an impractically large number of objects replaced by each participant (perhaps rivaling the number of cells recorded in our mice), the traditional noise ceiling might be more appropriate.

### Data Availability

All human navigation data generated by this study are publicly available at [insert link here]. All mouse CA1 data analyzed here are also publicly available and can be downloaded from [https://zenodo.org/records/14867736].

### Code Availability

All code necessary to reproduce this study are publicly available at [insert link here].

### Statistics

All statistical tests are noted as they appear in the text, and reported outcomes are uncorrected and two-tailed unless otherwise noted. In the case of nonparametric bootstrapped statistical tests, p-values reflect the uncorrected two-tailed probability that the compared populations differ. These were estimated by computing the measure of interest across 500 bootstrapped resamplings of each population with replacement and comparing these two distributions to one another nonparametrically. The p-value reflects the odds that these two distributions differ, to the resolution of these 250,000 comparisons. Box-and-whisker plots denote the minimum and maximum (whisker), the interquartile range (box), and median (line).

## Acknowledgements

This work was supported by generous startup funding from the University of Illinois Chicago as well as an LAS-CSSR Seed Grant from the University of Illinois Chicago. DGB, VHK, and MP were supported by CURA awards from the University of Illinois Chicago while contributing to this work. We are grateful to Russell A. Epstein for helpful feedback on an earlier draft of this work.

## Supplementary material

**Supplementary Figure 1.**
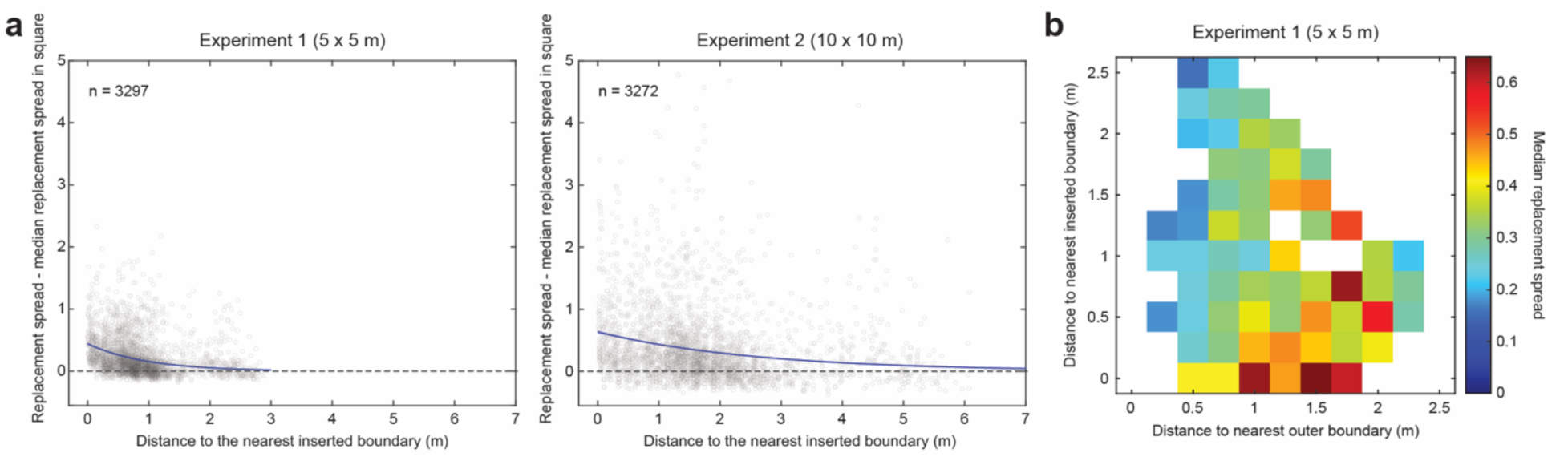
(**a**) Replacement spread as a function of distance from the nearest inserted boundary for accessible objects aggregated across all deformed configurations for experiments one and two. The best fitting functions are plotted here in purple, with 95^th^ percentile variability across 500 bootstraps plotted but difficult to see because the variability is minimal. The parameters yielding these fits are plotted in Figure 7e,f. (**b**) Median replacement spread for accessible objects as a function of distance to both the nearest inserted boundary and the nearest outer boundary of the environment for experiment one. Aggregated across all deformed configurations. Note that replacement spread generally increases as a function of distance to the nearest environmental boundary but decreases as a function of distance to the nearest inserted boundary. Bins with fewer than five replacements are excluded.

